# Characterization of full-length *CNBP* expanded alleles in myotonic dystrophy type 2 patients by Cas9-mediated enrichment and nanopore sequencing

**DOI:** 10.1101/2022.05.12.491603

**Authors:** Massimiliano Alfano, Luca De Antoni, Federica Centofanti, Virginia Veronica Visconti, Simone Maestri, Chiara Degli Esposti, Roberto Massa, Maria Rosaria D’Apice, Giuseppe Novelli, Massimo Delledonne, Annalisa Botta, Marzia Rossato

**Affiliations:** Department of Biotechnology, University of Verona, Strada Le Grazie 15, 37134 Verona, Italy; Department of Biomedicine and Prevention, Medical Genetics Section, University of Rome Tor Vergata, Via Montpellier 1, 00133 Rome, Italy; Department of Systems Medicine (Neurology), University of Rome Tor Vergata, Via Montpellier 1, 00133 Rome, Italy; Laboratory of Medical Genetics, Tor Vergata Hospital, viale Oxford 81, 00133 Rome, Italy; IRCCS Neuromed, Via Atinense, 18, 86077 Pozzilli, Molise, Italy; Department of Pharmacology, School of Medicine, University of Nevada, Reno, NV 89557, USA; Genartis s.r.l., Via P. Mascagni 98, 37060 Castel D’azzano (VR), Italy

**Keywords:** Myotonic dystrophy type 2, *CNBP* gene, repeat expansion, Cas9-mediated enrichment, nanopore sequencing

## Abstract

Myotonic dystrophy type 2 (DM2) is caused by CCTG repeat expansions in the *CNBP* gene, comprising 75 to >11,000 units and featuring extensive mosaicism, making it challenging to sequence fully-expanded alleles. To overcome these limitations, we used PCR-free Cas9-mediated nanopore sequencing to characterize *CNBP* repeat expansions at the single-nucleotide level in nine DM2 patients. The length of normal and expanded alleles can be assessed precisely using this strategy, agreeing with traditional methods, and revealing the degree of mosaicism. We also sequenced an entire ∼50-kbp expansion, which has not been achieved previously for DM2 or any other repeat-expansion disorders. Our approach precisely counted the repeats and identified the repeat pattern for both short interrupted and uninterrupted alleles. Interestingly, in the expanded alleles, only two DM2 samples featured the expected pure CCTG repeat pattern, while the other seven presented also TCTG blocks at the 3′ end, which have not been reported before in DM2 patients, but confirmed hereby with orthogonal methods. The demonstrated approach simultaneously determines repeat length, structure/motif and the extent of somatic mosaicism, promising to improve the molecular diagnosis of DM2 and achieve more accurate genotype– phenotype correlations for the better stratification of DM2 patients in clinical trials.

## INTRODUCTION

Myotonic dystrophy type 2 (DM2; MIM#602668) is an autosomal dominant multisystem disorder characterized by progressive proximal muscle weakness, myotonia, myalgia, calf hypertrophy, and multi-organ involvement with cataract, cardiac conduction defects, and endocrine disorders(Meola and Cardani 2015; Montagnese et al. 2017). The disease is caused by a (CCTG)_n_ repeat expansion in intron 1 of the *CNBP* gene (previously *ZNF9*; MIM*116955) on chromosome 3q21.3(Liquori et al. 2001). The CCTG repeat tract is part of a complex (TG)_v_ (TCTG)_w_ (CCTG)_x_ motif that, in healthy-range alleles, is generally interrupted by one or more GCTG, TCTG or ACTG (NCTG) motifs that confer repeat stability (Radvanszky et al. 2013) Mahyera et al., 2018) (Guo and Lam 2016). Non-pathogenic alleles contain up to 26 CCTG repeat units, whereas pre-mutations are composed of < 75 “pure” CCTG blocks whose clinical significance remains unclear(Mahyera et al. 2018)(Annalisa Botta et al. 2021). In DM2 patients, the number of repeats is between ∼75 and >11,000 units, among the largest reported so far in repeat-expansion disorders(Depienne and Mandel 2021). The DM2 mutation shows marked somatic instability and tends to increase in length over time within the same individual, but it does not show a strong bias toward intergenerational expansion, and genetic anticipation is rarely seen in DM2 families(Liquori et al. 2001),(Mahyera et al. 2018),(Kamsteeg et al. 2012),(Udd et al. 2003),(Thornton 2014).

The few genotype–phenotype studies reported so far in DM2 patients did not reveal any significant associations between the severity of the disease, including the age at onset, and the number of CCTG repeats(Day et al. 2003; Udd et al. 2003). The identification of such correlations is hindered by heterogeneity across tissues, somatic instability, and the technical challenges that must be overcome to measure repeat lengths accurately in such large microsatellite expansions. This has prevented the discovery of additional *in cis* genetic modifiers that may ameliorate or exacerbate DM2 disease symptoms. The genetic features of the *CNBP* microsatellite locus, as well as its extreme length and high CG content, have frustrated attempts to size and sequence expanded alleles in DM2 patients. Even the investigators in the original gene-discovery study were unable to sequence the entire CCTG array because of its length and the high level of somatic mosaicism(Liquori et al. 2001),(Bachinski et al. 2003),(Day et al. 2003).

According to international guidelines, current best practice for DM2 genetic testing relies mainly on PCR-based approaches(A Botta et al. 2006; Kamsteeg et al. 2012). These include an initial short-range PCR (SR-PCR) to exclude a DM2 diagnosis when two normal-range alleles are detected. When only one allele is visible, (CCTG)_n_ expansions can be identified by long-range PCR (LR-PCR) or quadruplet-repeat primed PCR (QP-PCR), leading to a ∼99% detection rate(Kamsteeg et al. 2012). However, neither method can define the exact length of large DM2 expansions, which requires the Southern blot analysis of digested genomic DNA and has a sensitivity of ∼80%(Day et al. 2003). The latter method is time consuming, requires large amounts of DNA, and is not included in the routine workflow of most diagnostic centers. Moreover, none of the methods described above can resolve to the single-nucleotide level and they have a limited ability to detect minor alleles and the degree of somatic mosaicism.

Third-generation long-read sequencing technologies provide an unprecedented opportunity to fully characterize DM2 expansions in terms of repeat size, allele configuration and base composition. Whereas most second-generation sequencing methods produce short reads, third-generation methods such as Oxford Nanopore Technologies (ONT) and PacBio SMRT sequencing facilitate the analysis of DNA fragments multiple kilobases in length, including large repetitive elements and their flanking regions. Furthermore, third-generation methods are generally based on PCR-free workflows, thus avoiding amplification-related biases(Hommelsheim et al. 2014) and challenges caused by regions with a high CG content (such as the *CNBP* locus). However, these technologies are more expensive and error-prone than first-generation and second-generation methods. Targeted approaches that maximize data production on a selected region of interest can compensate for such errors and provide cost-effective alternatives to whole-genome sequencing.

Targeted enrichment approaches coupled to long-read sequencing have already been used for the in-depth characterization of repeat expansions in fragile X syndrome (*FMR1*), Huntington’s disease (*HTT*) and neuronal intranuclear inclusion disease (*NOTCH2NLC*), although these expansions are significantly shorter than the (CCTG)_n_ repeats in DM2 patients(Dejesus-Hernandez et al. 2021; Ebbert et al. 2018; Giesselmann et al. 2019; Grosso et al. 2021; Hafford-Tear et al. 2019; Höijer et al. 2018; Mizuguchi et al. 2021; Sone et al. 2019; Tsai et al. 2017; Wallace et al. 2021; Wieben et al. 2019; Mitsuhashi and Matsumoto 2020). The longest allele thus far characterized by third-generation sequencing is a 21-kbp allele of the *C9orf72* gene that causes amyotrophic lateral sclerosis (ALS) and frontotemporal dementia (FTD) (Dejesus-Hernandez et al. 2021). In addition, many of the works cited above combined PacBio SMRT sequencing with LR-PCR(Mangin et al. 2021; Ciosi et al. 2021; Cumming et al. 2018), which is unsuitable for the analysis of DM2 expansions due to their extreme length and high GC content. Importantly, PacBio reads do not exceed 20 kbp in a PCR-free enrichment(Dejesus-Hernandez et al. 2021; Ebbert et al. 2018; Hafford-Tear et al. 2019; Höijer et al. 2018; Tsai et al. 2017; Wieben et al. 2019), and would not completely span DM2 pathogenic expansions (20 kbp on average, but up to 50 kbp).

To address these issues, we assessed the analysis of *CNBP* expansions using a combination of CRISPR/Cas9-based enrichment (Cas9-enrichment) and ONT sequencing, which can generate reads > 100 kbp in length(Payne et al. 2019; Iyer et al. 2022) and recently demonstrated valuable for the analysis of repetitive elements, e.g. telomeric and centromeric regions, in the completion of human genome (Sergey et al. 2022; Consortium 2022). In this manner, we sequenced full-length (CCTG)_n_ expansions in nine DM2 patients for the first time, including one mutated allele 47 kbp in length. Because this approach achieves single-nucleotide resolution, we were able to detect a previously unreported (TCTG)_n_ motif at the 3′ end of the *CNBP* expansion in seven of the DM2 patients. Our pilot study demonstrated that Cas9-mediated enrichment and long-read sequencing improves the DM2 diagnostic workflow, facilitating the in-depth characterization of *CNBP* expansions by accurately reporting the repeat length, structure/motif and degree of somatic mosaicism in a single analysis. In the future, this approach may enable more precise genotype–phenotype correlations and thus improve patient stratifications in clinical trials for personalized therapies.

## RESULTS

### Molecular characterization of DM2 patients using traditional methods

We analyzed nine DM2 patients (six males and three females, mean age = 46.4 ± 20 years) with existing molecular diagnoses based on a combination of PCR-based approaches (SR-PCR, LR-PCR and QP-PCR) to detect the presence of (CCTG)_n_ expansions in the *CNBP* gene (**Table 1**). Four patients were familial cases (A1–A4) whereas patients B, C, D, E and F were sporadic cases. The maximum length of the (CCTG)_n_ expansion could not be determined using routine diagnostic methods, so we digested genomic DNA and estimated the size of each allele by Southern blot analysis (**Figure 1A**). This suggested that the size of the microsatellite ranged from 20 to 48 kbp (**Figure S1** and **Table 1**). As expected, no signal was detected for healthy control subjects (CTR) or myotonic dystrophy type 1 (DM1) patients (**Figure S1**). The characterization of normal *CNBP* alleles by SR-PCR and Sanger sequencing revealed the presence of eight short interrupted alleles with the structure (TG)_17-24_ (TCTG)_6-9_ (CCTG)_5-7_ GCTG CCTG TCTG (CCTG)_7_ and one short uninterrupted allele with the structure (TG)_19_ (TCTG)_9_ (CCTG)_12_, matching our previous results for an Italian population(Annalisa Botta et al. 2021) (**Table 1**).

**Table 1.**
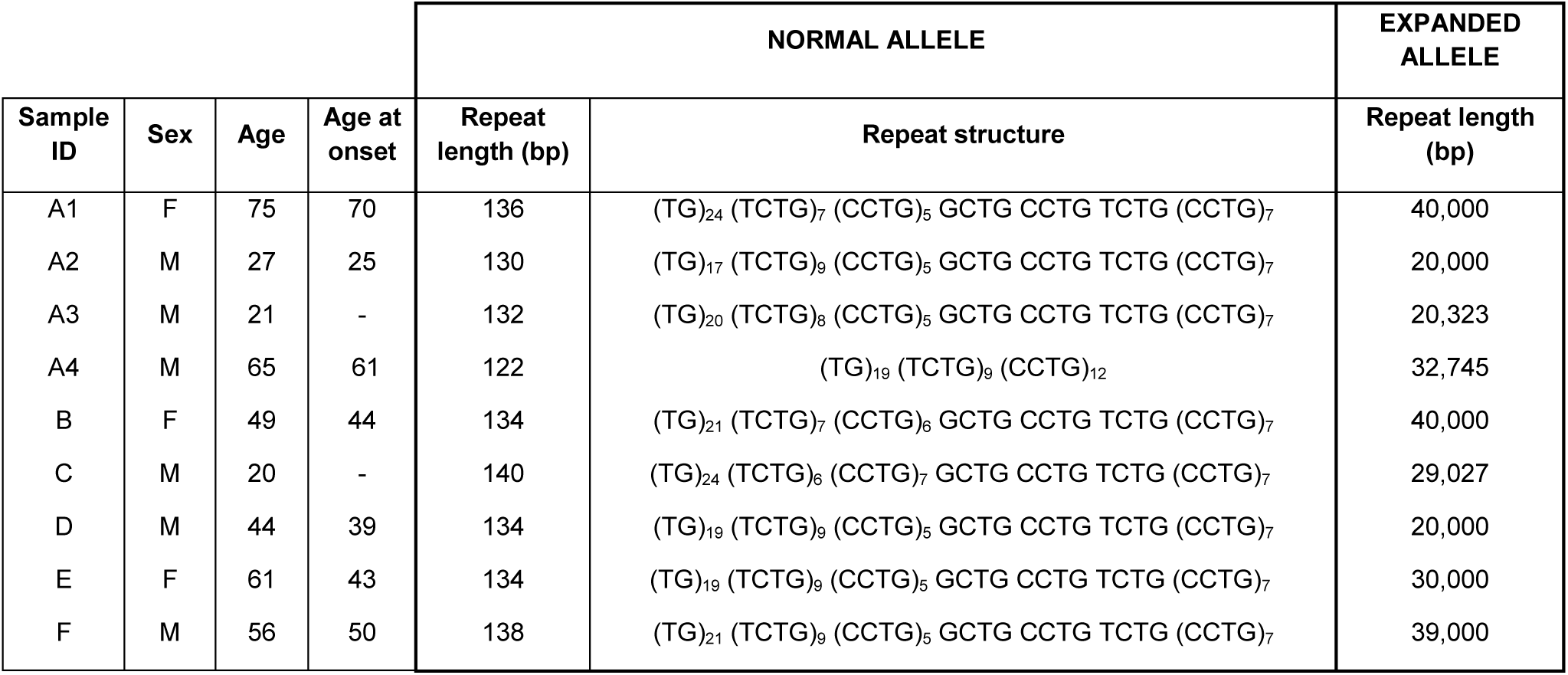
Demographic and molecular features of the DM2 patients. For each patient, the table shows the ID, sex, age, age at onset, repeat length and structure of the *CNBP* normal allele characterized by Sanger sequencing, and the estimated repeat length of the expanded allele determined by Southern blotting (–, not available).

**Figure 1.**
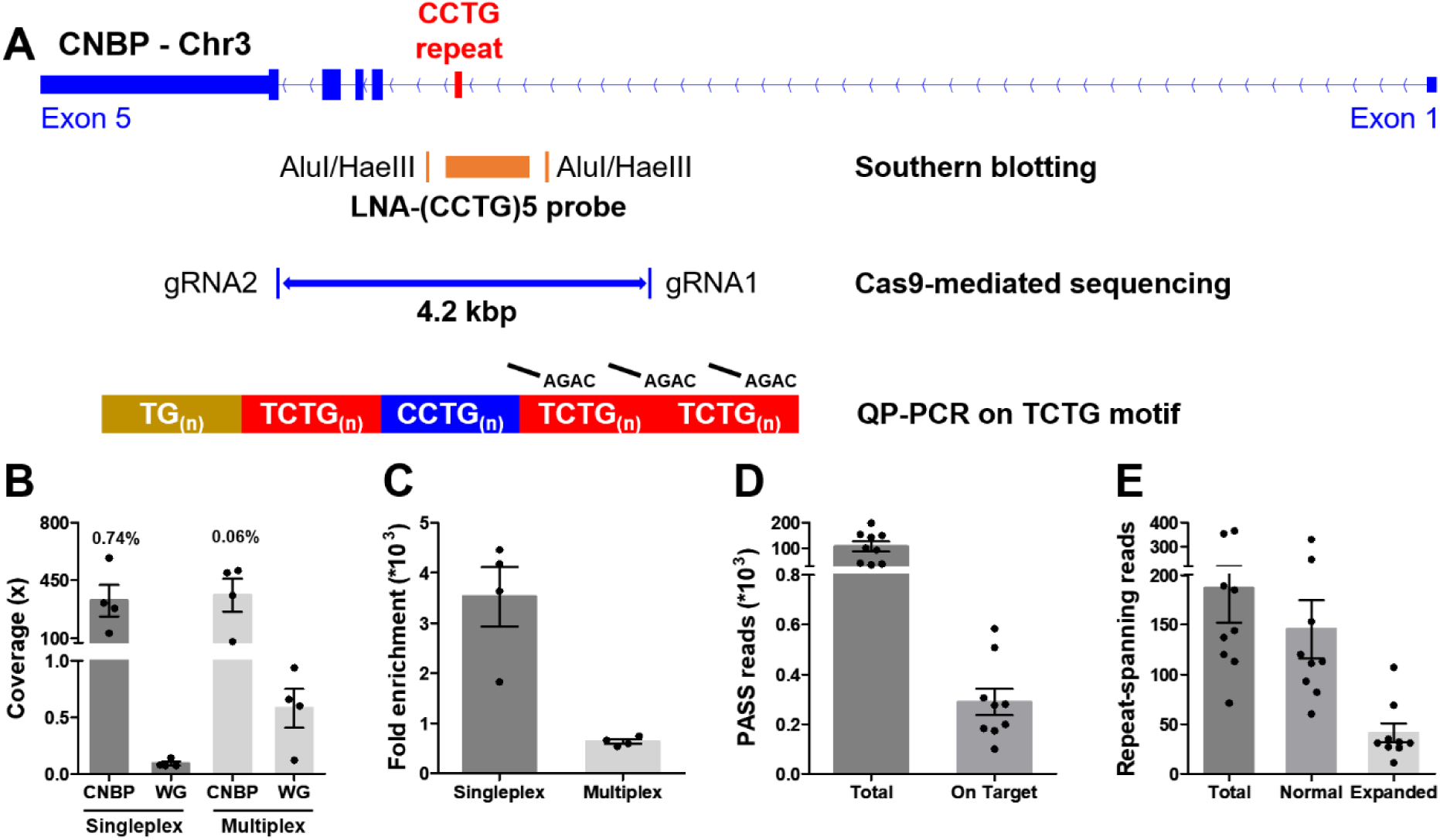
Cas9-mediated sequencing of the *CNBP* microsatellite. **(A)** Experimental methods applied retrospectively to study the *CNBP* microsatellite in nine confirmed DM2 patients. The positions of AluI and HaeIII restriction sites (142 bp upstream and 108 bp downstream of the CCTG array, respectively) and the (CCTG)_5_ probe (orange rectangle) for Southern blot hybridization are shown, along with the gRNAs cleavage site for Cas9-mediated enrichment (boundaries of blue line = 4.2 kb) and the annealing position of P4TCTG primers for QP-PCR. **(B)** Average coverage and **(C)** fold-enrichment obtained by Cas9-mediated enrichment in DM2 patients following singleplex (n = 4) or multiplex (n = 4) runs. Numbers above bars represent the percentage of on-target reads. **(D)** Total and on-target number of PASS reads generated for each DM2 patient and **(E)** number of reads fully spanning the *CNBP* microsatellite and attributable to either the normal or expanded alleles. Data are plotted as means ± SEM. WG, whole genome.

### *CNBP* repeat expansion analysis by Cas9-mediated enrichment coupled to ONT sequencing

We characterized the full-length *CNBP* expansions at single-nucleotide resolution by ONT sequencing following Cas9-mediated enrichment. Accordingly, we designed two gRNAs to excise a 4.2-kbp fragment spanning the *CNBP* repeat on chromosome 3q21.3 (**Figure 1A and Table S1**).

Genomic DNA from the nine DM2 patients was analyzed in four singleplex and four multiplex runs, the latter applied to clinical samples here for the first time (**Table S1**). Cas9-mediated sequencing achieved good target coverage in all experiments, with 346 ± 64 reads (mean ± SEM) on the *CNBP* locus (**Figure 1B and Table S1**). Singleplex runs has consistently lower background than multiplex runs (0.08× vs 0.57×), and thus achieved a higher average fold enrichment (3,521-fold vs 637-fold) (**Figure 1B-C and Table S1**). Collectively, for each DM2 patient, we generated a mean total of 105,737 PASS reads, 308 of which were on target (**Figure 1D**) and 186 of which completely spanned the normal or expanded alleles (**Figure 1E**). Only these “complete sequences” were used for subsequent analysis, representing ∼78% and ∼22% of the normal and expanded repeat-spanning reads, respectively (**Figure 1E**).

The *de novo* assembly of reads derived from the normal *CNBP* alleles in DM2 patients (145 on average per sample, IQR = 67; **Figure 1E**) showed that the complex (TG)_v_ (TCTG)_w_ (CCTG)_x_ (NCTG)_y_ (CCTG)_z_ repeat ranged in size from 122 to 141 bp, corresponding to 12–15 CCTG quadruplets (**Table 2** and **Figure 2B**). The size and repeat pattern in each patient were largely consistent with the Sanger sequencing data (99.5% mean accuracy, Pearson’s r = 0.971, p < 0.0001; **Figure 2C**), with six patients showing a perfect match, two differing at a single nucleotide position and only one differing at two nucleotide positions (**Table 2**).

**Table 2.** *CNBP* repeat analysis based on Cas9-mediated sequencing. For each patient, the table shows the characteristics of normal and expanded *CNBP* alleles based on the analysis of ONT sequencing data, in terms of length and structure. For the normal alleles, the table reports the percentage identity of consensus sequences reconstructed from ONT or Sanger sequencing data and the incongruences in ONT-derived sequences are highlighted in bold. For the expanded alleles, the table indicates the proportion of reads carrying the unexpected TCTG motif at the 3′ end of the *CNBP* microsatellite expansion.

| NORMAL ALLELE | Sample ID | Repeat length (bp) | Repeat structure | Identity with Sanger sequence |
| --- | --- | --- | --- | --- |
|  | A1 | 136 | (TG)<sub>24</sub> (TCTG)<sub>7</sub> (CCTG)<sub>5</sub> GCTG CCTG TCTG (CCTG)<sub>7</sub> | 100.0% |
|  | A2 | 131 | (TG)<sub>17</sub> TGCTG (TCTG)<sub>8</sub> (CCTG)<sub>5</sub> GCTG CCTG TCTG (CCTG)<sub>7</sub> | 99.2% |
|  | A3 | 132 | (TG)<sub>20</sub> (TCTG)<sub>8</sub> (CCTG)<sub>5</sub> GCTG CCTG TCTG (CCTG)<sub>7</sub> | 100.0% |
|  | A4 | 122 | (TG)<sub>19</sub> (TCTG)<sub>9</sub> (CCTG)<sub>12</sub> | 100.0% |
|  | B | 134 | (TG)<sub>21</sub> (TCTG)<sub>7</sub> (CCTG)<sub>6</sub> GCTG CCTG TCTG (CCTG)<sub>7</sub> | 100.0% |

**Table 2. CNBP repeat analysis based on Cas9-mediated sequencing.**
|  | C | 141 | (TG)<sub>24</sub> **TGCTG** (TCTG)<sub>5</sub> (CCTG)<sub>7</sub> GCTG CCTG TCTG (CCTG)<sub>7</sub> | 99.3% |
| --- | --- | --- | --- | --- |
|  | D | 138 | **(TG)**<sub>21</sub> (TCTG)<sub>9</sub> (CCTG)<sub>5</sub> GCTG CCTG TCTG (CCTG)<sub>7</sub> | 97.1% |
|  | E | 134 | (TG)<sub>19</sub> (TCTG)<sub>9</sub> (CCTG)<sub>5</sub> GCTG CCTG TCTG (CCTG)<sub>7</sub> | 100.0% |
|  | F | 138 | (TG)<sub>21</sub> (TCTG)<sub>9</sub> (CCTG)<sub>5</sub> GCTG CCTG TCTG (CCTG)<sub>7</sub> | 100.0% |
| EXPANDED ALLELE | Sample ID | Repeat length (bp)<br>(min - max) | Repeat structure | Number of reads carrying the TCTG motif |
|  | A1 | 3241 - 46685 | (TG)<sub>20</sub> (TCTG)<sub>7</sub> (CCTG)<sub>1000-12000</sub> (TCTG)<sub>0-10</sub> | 14 (45%) |
|  | A2 | 864 - 23779 | (TG)<sub>18</sub> (TCTG)<sub>7</sub> (CCTG)<sub>1000-4500</sub> (TCTG)<sub>0-2000</sub> | 92 (86%) |
|  | A3 | 4429 - 18983 | (TG)<sub>19</sub> (TCTG)<sub>7</sub> (CCTG)<sub>3000-5000</sub> (TCTG)<sub>0-25</sub> | 8 (73%) |
|  | A4 | 660 - 34284 | (TG)<sub>18</sub> (TCTG)<sub>7</sub> (CCTG)<sub>250-8000</sub> (TCTG)<sub>0-1500</sub> | 9 (29%) |
|  | B | 344 - 23358 | (TG)<sub>18</sub> (TCTG)<sub>7</sub> (CCTG)<sub>300-4000</sub> (TCTG)<sub>0-400</sub> | 6 (23%) |
|  | C | 700 - 31753 | (TG)<sub>20</sub> (TCTG)<sub>7</sub> (CCTG)<sub>150-8000</sub> | 0 (0%) |
|  | D | 383 - 19143 | (TG)<sub>18</sub> (TCTG)<sub>7</sub> (CCTG)<sub>100-4000</sub> (TCTG)<sub>0-1000</sub> | 3 (11%) |
|  | E | 848 - 25162 | (TG)<sub>18</sub> (TCTG)<sub>6</sub> (CCTG)<sub>200-6200</sub> | 0 (0%) |
|  | F | 1533 - 32824 | (TG)<sub>15</sub> (TCTG)<sub>10</sub> (CCTG)<sub>400-6000</sub> (TCTG)<sub>0-2000</sub> | 15 (43%) |

**Figure 2.**
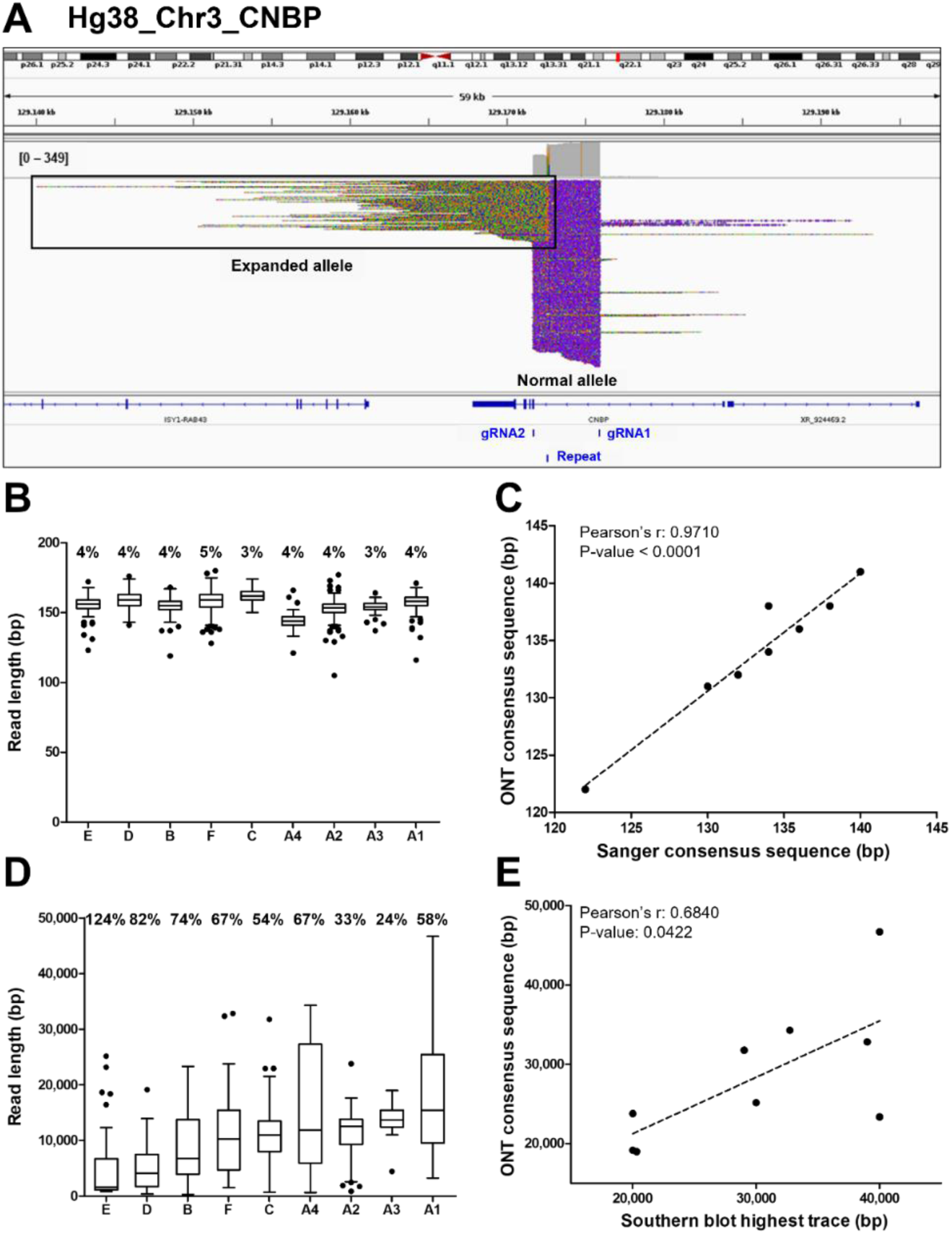
Analysis of ONT sequencing data from normal and expanded *CNBP* alleles. **(A)** Integrative Genomics Viewer (IGV) visualization of ONT sequencing data at the *CNBP* locus of a representative DM2 patient following Cas9-mediated enrichment. Reads generated from the normal allele feature clear cuts on both sides of the *CNBP* repeat, whereas those derived from the expanded allele are longer, soft-clipped and do not match the reference genome, as expected. Length distributions of reads derived from the normal alleles **(B)** and expanded alleles **(D)** of each patient. Boxes represent the interquartile range (IQR) of lengths, the horizontal line is the median, whiskers and outliers are plotted according to Tukey’s method. **(C)** Correlation between the length of ONT and Sanger consensus sequences for the normal allele (n = 9). **(E)** Correlations between the maximum length of ONT sequences (longest complete read) and the upper edge of the Southern blot trace for the expanded allele (n = 9). Numbers on top of panels **(B)** and **(D)** indicate the coefficient of variation of normal and expanded alleles, respectively.

Reads derived from the expanded alleles (41 on average per sample, IQR = 11; **Figure 1E**) ranged from 344 bp to as much as 46.6 kbp (**Figure 2D**), confirming the presence of extremely large expansions in these patients. To our knowledge, the latter is the longest repeat expansion analyzed thus far at single-nucleotide resolution(Mizuguchi et al. 2021; Sone et al. 2019; Giesselmann et al. 2019; Wallace et al. 2021) and is one of the longest DNA fragments captured by Cas9-mediated enrichment with no specific adjustment(Gilpatrick et al. 2020; Iyer et al. 2020). Considering average values per sample, the number of repetitive quadruplets varied from 1,371 to 4,421, corresponding to expansion lengths of 5,485–17,685 bp. Moreover, in each patient, the longest molecule sequenced with ONT (derived from the expanded allele) largely agreed with the Southern blot estimates (Pearson’s r = 0.6840, p = 0.0422, **Figure 2E**), with the exception of sample (B). Even within the same individual, the mean size of reads derived from the expanded allele was consistently more variable than that from the normal allele (65% ± 0.091 vs 4% ± 0.002; **Figure 2B-D, Figure 3 and Figure S2**). Such pronounced variability within each DM2 patient indicates extensive mosaicism, in agreement with previous reports(Liquori et al. 2001; Bachinski et al. 2003; Day et al. 2003).

**Figure 3.**
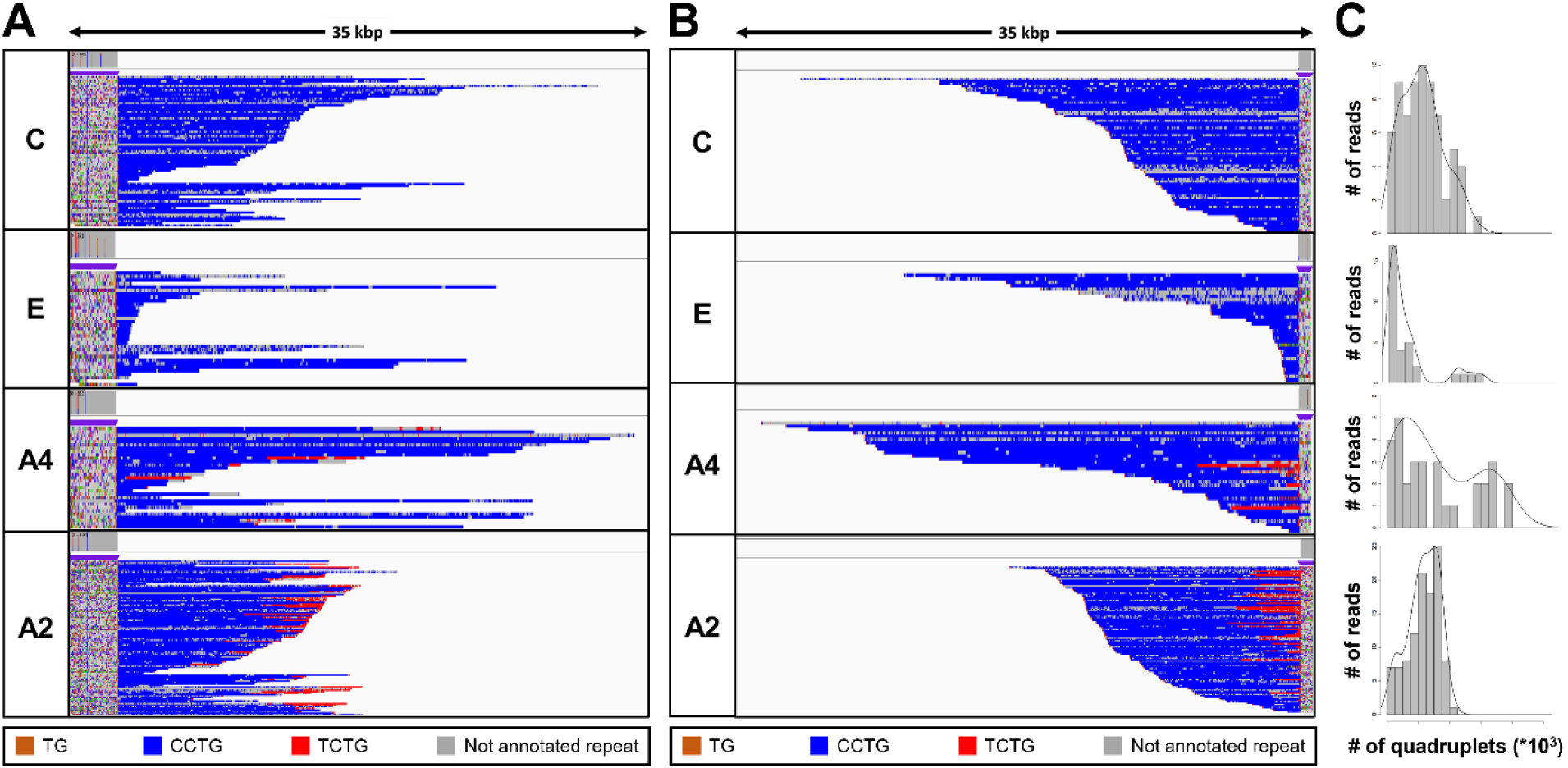
Analysis of the expanded-repeat *CNBP* alleles in DM2 patients. Integrative Genomics Viewer (IGV) visualization (35-kbp windows) of ONT targeted sequencing data from the expanded alleles of four representative DM2 patients. Complete reads were aligned at the 3′ end **(A)** and then at the 5′ end **(B)** in order to identify the repeat pattern that characterizes the expanded microsatellite locus. Each motif in the expanded alleles was visualized using a different color, as indicated in the key. Samples C and E contained a “pure” CCTG expansion (blue) whereas samples A4 and A2 also contained the unexpected TCTG motif (red) downstream of CCTG. (**C**) Abundance of quadruplets identified in each patient. The *y*-axis shows the number of ONT reads with a certain number of repeats, whereas the *x*-axis shows the number of quadruplet repeats identified. ONT reads were grouped into 500-bp bins. The gray line represents the estimated kernel density of the underlying solid gray distribution of ONT reads.

To characterize the repeat pattern across the expanded microsatellite locus, we identified the quadruplet motifs in each individual read, and highlighted them with distinct colors after aligning “complete sequences” at the 5′ and 3′ ends (**Figure 3 and Figure S2**). The total number of quadruplets was highly variable, thus confirming the extensive donor-dependent mosaicism described above (**Figure 3C and Figure S2C**). The uninterrupted (TG)_v_ (TCTG)_w_ (CCTG)_x_ motif that characterizes expanded *CNBP* alleles was found at the 5′ end of the repeat locus in all nine patients (**Figure 3A, Figure S2A and Table 2**). However, only two patients featured a “pure” pattern of (CCTG)_n_ repeats. In the remaining seven, we observed additional (TCTG)_n_ arrays (colored in red) at the 3′ end of the CCTG expansion, which has been never reported in DM2 patients before (**Figure 3B, Figure S2B and Table 2**). When present, the TCTG motif was detected in a highly variable fraction of sequences (11–86% of the expanded allele reads, **Table 2**), and differed widely in length (40–8,000 bp) between donors and within the same individual (**Figure 3, Figure S2 and Table 2**).

### Analysis of the TCTG repeat using orthogonal methods

To confirm the presence of the (TCTG)_n_ motifs in the *CNBP* expanded alleles, we used a traditional QP-PCR method for the selective amplification of TCTG blocks at the 3′ end of the (CCTG)_n_ array (**Figure 1A**). In agreement with the ONT data, QP-PCR analysis using primer P4TCTG revealed an electrophoretic profile compatible with the presence of (TCTG)_n_ downstream of the (CCTG)_n_ expansion in seven DM2 samples (**Figure 4A**). The intensity and pattern of fluorescent peaks obtained using primer P4TCTG was more variable across samples compared to the routine protocol using the standard P4CCTG primer, possibly due to different levels of somatic mosaicism in the TCTG and CCTG expansions. A 260-bp signal was visible in all samples, including the DM2 negative controls (one DM1 positive patient and one healthy subject CTR), suggesting it was a PCR artifact (**Figure 4**). Interestingly, the two patients with “pure” (CCTG)_n_ expansions based on ONT data (patients C and E) did not yield amplification peaks within the expansion range with the P4TCTG primer (**Figure 4B**). Fluorescent signals < 140 bp were visible in these samples because normal *CNBP* alleles also contain (TCTG)_n_ repeats at their 5′ end (**Table 1**). The direct sequencing of QP-PCR products generated using primer P4TCTG confirmed the presence of (TCTG)_n_ motif in DM2 patients A1, A2, A3, A4, B, D and F, thus supporting the ONT data (**Figure 4 and Figure S3)**.

**Figure 4.**
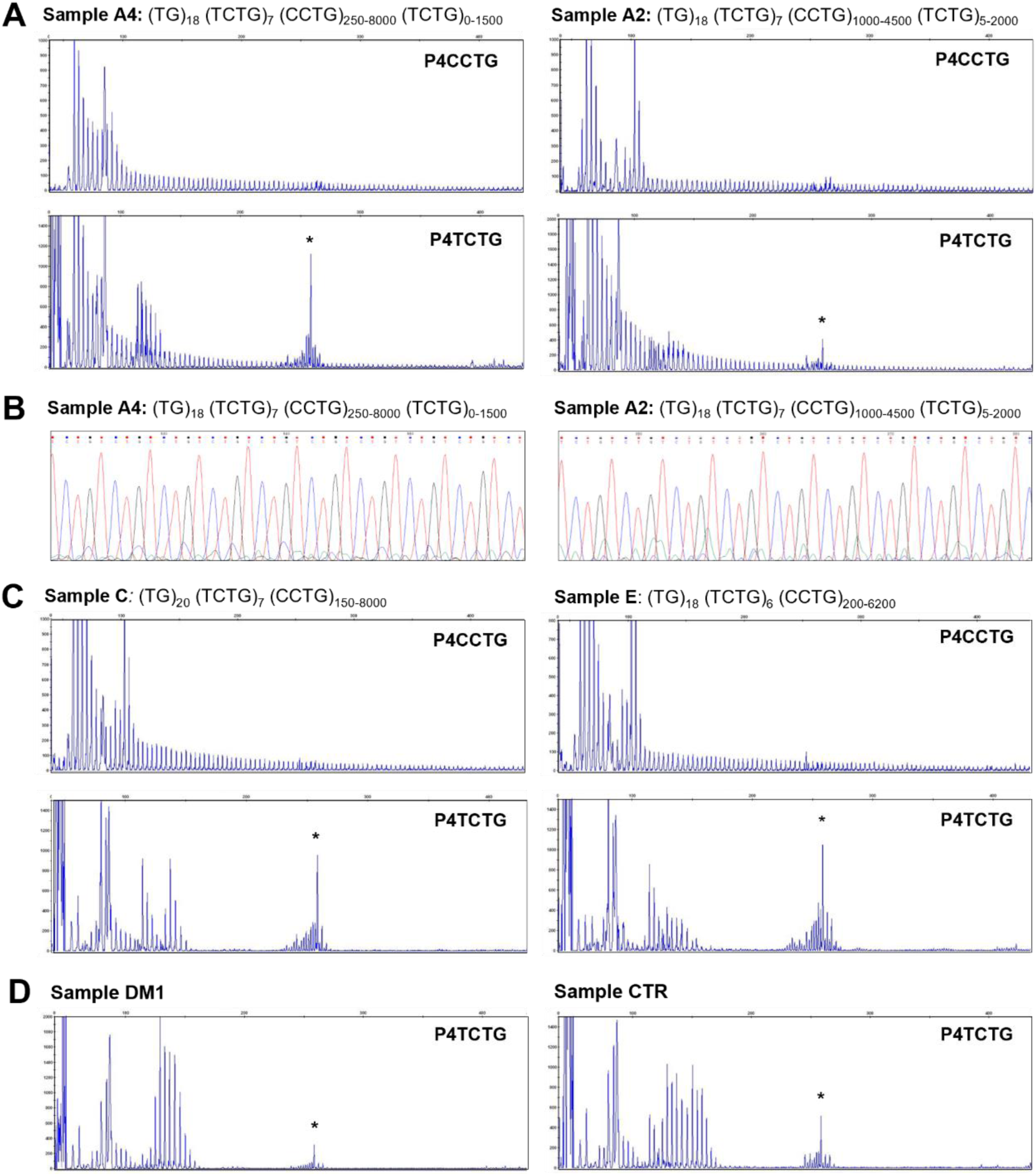
Analysis of the TCTG motif by QP-PCR and Sanger sequencing. (**A**) Representative QP-PCR profiles of genomic DNA samples from patients A2 and A4 and showing the presence of the TCTG block. Upper panels show QP-PCR results using the conventional P4CCTG primer. Lower panels show QP-PCR results using primer P4TCTG. (**B**) Sanger sequencing of the QP-PCR products from P4TCTG reaction confirming the presence of the TCTG sequence. (**C**) QP-PCR profiles of genomic DNA samples from patients C and E showing only the traditional DM2 motif CCTG. For each patient, the composition of *CNBP* expanded alleles above the QP-PCR tracks reflects the ONT sequencing data. Asterisks (*) indicate non-specific signals, also visible in the QP-PCR profiles of DM1 and CTR samples **(D)**.

## DISCUSSION

The analysis of extremely long microsatellite expansions is challenging, preventing the in-depth characterization of *CNBP* mutations underlying DM2 and the identification of genotype–phenotype correlations in these patients. We have demonstrated for the first time the use of Cas9-mediated enrichment and ONT long-read sequencing to analyze the full (CCTG)_n_ expansion in DM2 patients at single-nucleotide resolution. We were able to characterize normal and expanded *CNBP* alleles, simultaneously revealing the repeat length, structure/motif and extent of somatic mosaicism, which is not possible with traditional methods, even if several are used in combination.

The paucity of genotype–phenotype data for DM2 reflects the historical inability to determine the size of CCTG repeats, especially in the largest expansions, which traditionally was based on the Southern blot analysis of genomic DNA. This labor-intensive and time-consuming technique is becoming obsolete because it requires large amounts of high-molecular-weight DNA. LR-PCR can be used instead, but the performance of this technique is poor in regions with a high CG content, and it cannot accommodate the very large expanded alleles (> 15 kbp) often found in DM2. Therefore, although LR-PCR achieves the sensitive detection of DM2 mutations, the full length of the (CCTG)_n_ expansion cannot be determined in all patients. Our Cas9-targeted sequencing protocol overcomes these limits by focusing long-read sequencing data on the *CNBP* microsatellite, with reads spanning the entire expansion. The length of normal and expanded alleles in DM2 patients determined using this new approach closely matched the values obtained with the traditional reference methods. The mean size of normal *CNBP* alleles was 134 bp whereas expanded alleles ranged from 344 to 46,685 bp, in agreement with previous reports(Meola and Cardani 2015; Montagnese et al. 2017; Annalisa Botta et al. 2021). A single incongruence was observed for one expanded allele in patient B, where ONT sequencing underestimated the size determined by Southern blotting (∼20 *vs* ∼40 kbp). A possible explanation is the presence of damaged DNA in the sample, reflecting its long-term storage in a biobank. Southern blotting involves the fractionation of double-stranded DNA, which would be unaffected by the presence of nicked strands, whereas ONT sequencing involves the analysis of single DNA strands, so the presence of nicks would have a profound effect (Oxford Nanopore Community 2021). Even so, Cas9-mediated enrichment allowed us to sequence DM2 alleles up to ∼50 kbp in length at single-nucleotide resolution, which has not been reported previously for DM2 or any other repeat-expansion disorder using this approach (Giesselmann et al. 2019; Mizuguchi et al. 2021; Sone et al. 2019; Wallace et al. 2021; Gilpatrick et al. 2020; Iyer et al. 2020). The analysis of such long and repetitive alleles required the coupling of a PCR-free enrichment protocol to ONT sequencing because even other long-read sequencing technologies cannot accommodate this read length in targeted sequencing experiments. For example, PacBio long-read sequencing was previously used to sequence repeat expansions in DM1, which is also characterized by long alleles of 4–6 kbp, but the microsatellites were first amplified by PCR(Mangin et al. 2021; Cumming et al. 2018). Even when coupled to PCR-free enrichment approach based on Cas9, the length of PacBio sequencing reads cannot exceed 20 kbp(Dejesus-Hernandez et al. 2021; Ebbert et al. 2018; Hafford-Tear et al. 2019; Höijer et al. 2018; Tsai et al. 2017; Wieben et al. 2019).

From a technical perspective, we achieved > 500× enrichment on *CNBP* using the Cas9 protocol, which is robust and comparable to similar assays for the assessment of microsatellite length(Giesselmann et al. 2019; Mizuguchi et al. 2021; Sone et al. 2019; Wallace et al. 2021). We also compared Cas9-mediated singleplex *vs* multiplex enrichment protocols for the first time on clinical samples and observed a consistently lower performance (10-fold lower enrichment, with 70% unclassified reads) in the multiplex environment, as reported by other ONT users in the ONT for other type of samples (Oxford Nanopore Community 2022). Further improvements are therefore required before the multiplexing protocol is suitable, for example by combining the Cas9 protocol with the recently introduced “Read Until” feature of ONT sequencing to exclude the background noise. This allows selective sequencing of pre-defined target DNA molecules and has been used to characterize pathogenic repeats(Stevanovski et al. 2021). Alternatively, costs could be optimized by analyzing single samples using Flongles, which produce less sequencing data than regular ONT flow cells but reduce costs by 90%.

The bioinformatic pipeline we used to analyze the *CNBP* microsatellite sequences also allowed us to recognize the repeat pattern and precisely count the repeats, including the short interrupted and uninterrupted alleles typifying the Italian population(Annalisa Botta et al. 2021). Consensus sequences generated from the normal allele shared a high degree of identity (99.5%) and significant correlation with state-of-the-art Sanger sequences, aside from a few nucleotide positions. These inconsistences probably reflect ONT sequencing errors and could be addressed by using the most recent base-calling algorithm and eventually the more accurate Q20+ chemistry.

In the expanded alleles of seven DM2 patients, the anticipated (CCTG)_n_ repeat was accompanied by a previously unreported (TCTG)_n_ repeat located at the 3′ end of the (CCTG)_n_ array. When present, the atypical motif varied in length between donors and within each sample (40–8000 bp). Repeat interruptions within the expanded array have been reported in 3–8% of DM1 patients(Tomé et al. 2018; Braida et al. 2010; M Santoro et al. 2017; Massimo Santoro et al. 2013; Ballester-Lopez et al. 2020; Pešović et al. 2018; Miller et al. 2020; Radvansky et al. 2011; De Siena et al. 2018; Annalisa Botta et al. 2017; Massimo Santoro et al. 2015; Addis et al. 2012; Fontana et al. 2020; Lian et al. 2016; Leeflang and Arnheim 1995; Musova et al. 2009; Cumming et al. 2018), but have not been described in DM2 before. This may reflect the challenge of sequencing complete *CNBP* expanded alleles and/or the use of a primer containing “pure” (CCTG)_n_ repeats for diagnostic QP-PCR. Given that (TCTG)_n_ repeats were present in a highly variable proportion of expanded alleles (11–86%), always in the presence of the typical (CCTG)_n_ repeats, technical bias may have revealed only the “pure” (CCTG)_n_ repeats. Indeed, a modified QP-PCR protocol using a primer containing five TCTG units – (TCTG)_5_T – was able to confirm the ONT sequencing data. From a biological perspective, the (TCTG)_n_ motif may have arisen through DNA duplication/repair errors or spontaneous DNA damage in the somatic cells of DM2 patients. Although the presence of this motif may be biologically relevant in the context of DM2, our data must be interpreted with caution. First, the motif was discovered in a small set of patients, most belonging to the same family, so confirmation requires a larger DM2 cohort enrolled in a multi-center international study. Second, considering the known limitations of ONT sequencing when presented with low-complexity regions such as homopolymers, the length and recurrence of such motifs should be investigated using other long-read methods when they are sufficiently advanced to sequence the expanded alleles completely.

The Cas9-targeted sequencing approach also allowed us to estimate the degree of somatic mosaicism for the mutated alleles, either “pure” or “interrupted”. On average, the allele length within each patient varied by 65%. Mosaicism plays an important role in the development of disease symptoms, so establishing the relative proportion of expanded alleles in the lower and upper mutation range could add prognostic value, significantly improving genetic counselling for DM2. Extreme mosaicism (more than expected based on previous studies) has also been detected when long-read sequencing is applied to other repeat-expansion disorders, suggesting that such techniques achieve higher resolution(Loomis et al. 2013; Mizuguchi et al. 2021; Mangin et al. 2021). As already demonstrated for DM1(Cumming et al. 2019; Monckton et al. 1995), the progenitor allele length (i.e., the length of the CCTG repeat transmitted by the affected parent) is one genetic determinant that influences the age at onset of DM2 symptoms, and that age is further modified by individual-specific differences in the level of somatic instability. Notably, our method accurately distinguished between the shortest expanded allele and the normal allele.

Another advantage of the approach demonstrated hereby is that PCR-free analysis potentially allows the direct assessment of DNA methylation, as already reported for other repeat-linked diseases(Fukuda et al. 2021; Giesselmann et al. 2019). This can provide additional information concerning the impact of expansions on the functionality of the *CNBP* gene. The methylation of the *CNBP* gene has been analyzed using a pyrosequencing method, revealing hypomethylation of CpG sites upstream and hypermethylation of CpG sites downstream of the (CCTG)_n_ expansion in DM2 patients and healthy individuals, with no significant differences between these groups(Massimo Santoro et al. 2018). However, it remains possible that the DM2 mutation could have epigenetic effects in other regulatory regions of the *CNBP* gene and/or in different tissues.

Given the ability of our method to simultaneously determine the size, single-nucleotide composition and degree of somatic mosaicism of DM2 repeat expansions, all of which are prerequisites for genotype–phenotype correlation, ONT sequencing could be included in the DM2 diagnostic workflow to improve the information content available for genetic counseling. Subsequent investigations using larger patient cohorts will be necessary to unravel the impact of each of these genetic factors on the onset and severity of the disease.

Taken together, our pilot study has demonstrated the potential of PCR-free long-read sequencing for the genetic assessment of DM2, allowing us to investigate both the length and genetic features of normal and expanded alleles in a single round of analysis. The use of such an approach in larger cohorts will increase the accuracy of genotype–phenotype correlations and enhance the information content available for DM2 genetic counselling.

## MATERIALS AND METHODS

### DM2 patients

We retrospectively analyzed nine genetically confirmed DM2 Italian patients, whose enrollment in the study was approved by the institutional review board of Policlinico Tor Vergata (document no. 232/19). All experimental procedures were carried out according to The Code of Ethics of the World Medical Association (Declaration of Helsinki). Informed consent was obtained from all nine participants and all samples and clinical information were anonymized immediately after collection using a unique alphanumeric identification code. Sociodemographic data for the DM2 patients are summarized in **Table 1**.

### Southern blotting

Genomic DNA was extracted from 500 µL of anti-coagulated peripheral blood using a Flexigene DNA Kit (Qiagen, Hilden, Germany) and diluted to a final volume of 25 μl with double-distilled water. The quality and quantity of DNA was assessed using a Denovix spectrophotometer and by 1% agarose gel electrophoresis. *CNBP* expanded alleles were detected as previously described(Nakamori et al. 2009), with modifications. Briefly, 2 μg of genomic DNA was digested with AluI and HaeIII and the fragments were resolved by 0.4% agarose gel electrophoresis at 40 V for 40 h. After denaturation and neutralization, the DNA was transferred to a nylon membrane (MilliporeSigma, Burlington, Massachusetts) and fixed by UV cross-linking using a Stratalinker 2400 (Stratagene, San Diego, California). The membrane was hybridized for 16 h at 65 °C with a digoxigenin (DIG)-labeled locked nucleic acid (LNA) probe (CCTG)_5_ at a concentration of 10 pmol/mL. After washing at high stringency, the signal was revealed using the DIG High Prime DNA Labeling and Detection Starter Kit II (Roche, Basel, Switzerland) and visualized using an ImageQuant LAS 4000 device (GE Healthcare, Chicago, Ilinois). Bands were sized by running two sets of molecular weight markers alongside the samples: DNA Molecular Weight Marker XV (Expand DNA Molecular Weight Marker, Roche) and λ DNA-HindIII Digest (New England Biolabs, Ipswich, Massachusetts).

### SR-PCR, QP-PCR and Sanger sequencing

SR-PCR products were generated as reported earlier(Kamsteeg et al. 2012,(A Botta et al. 2006). QP-PCR targeting the 3′ end of the (CCTG)_n_ repeat array was carried out as previously described(Catalli et al. 2010; Musova et al. 2009), with modifications. Specifically, the repeat primer P4TCTG-agc gga taa caa ttt cac aca gga TCT GTC TGT CTG TCT GTC TGT (lower case letters indicate the primer tail that does not complement the repeat) was combined with primers CL3N58_DR-[FAM]-GCC TAG GGG ACA AAG TGA GA and P3-AGC GGA TAA CAA TTT CAC ACA GGA to target the most 3′ (TCTG)_n_ interruptions. The length of the *CNBP* unexpanded alleles and QP-PCR products were determined by capillary electrophoresis on the 3500 Genetic Analyzer followed by analysis using GeneMapper 6 (Applied Biosystems, Waltham, Massachusetts). The SR-PCR and QP-PCR products were purified using the ExoSAP protocol, directly sequenced using the Big Dye Terminator Cycle Sequencing Kit v3.1 (Thermo Fisher Scientific, Waltham, Massachusetts) and visualized by capillary electrophoresis on the 3500 Genetic Analyzer as above.

### Cas9-mediated enrichment coupled to ONT sequencing

For DM2 patients A1–A4, B and C, genomic DNA was extracted from 0.2–0.5 mL of peripheral blood using the Nanobind CBB Big DNA HMW Kit (Circulomics, Baltimore, Maryland). For DM2 patients D, E and F, genomic DNA was extracted from 1–2 mL whole blood using the NucleoSpin Blood L Kit (Macherey-Nagel, Düren, Germany). Regardless of the extraction method, the DNA was resuspended in Tris-EDTA buffer (pH 8.0) and the quantity was determined using a Qubit fluorometer (Thermo Fisher Scientific) and Qubit dsDNA BR Assay Kit (Thermo Fisher Scientific). DNA integrity was assessed using a TapeStation 4150 device, Genomic DNA ScreenTape and Genomic DNA Reagents (ladder and sample buffer) all from Agilent Technologies (Santa Clara, California).

We designed crRNAs using the online tool CHOPCHOP (https://chopchop.cbu.uib.no/) following ONT’s recommendations (https://community.nanoporetech.com/info_sheets/targeted-amplification-free-dna-sequencing-using-crispr-cas/v/eci_s1014_v1_reve_11dec2018), and making sure that the excised fragment was at least 3 kbp in length. Candidate crRNAs were manually checked for unique mapping by aligning them to the human genome (Hg38) using BLAST and excluding regions overlapping common single-nucleotide polymorphisms (MAF > 0.01, dbSNP database). The final crRNAs were prepared by Integrated DNA Technologies (Coralville, Iowa): 5′-CCA CCT GAT TCA CTG CGA TA-3′ with genomic coordinates Chr3:129,175,929-129,175,948, and 5′-GGC TTC TCA TTC CAC GAC CA-3′ with genomic coordinates Chr3:129,171,664-129,171,683. The reaction mixture comprised 10 µM of each crRNA, 10 µM of transactivation crRNA (tracrRNA) and 62 µM Alt-R S.p. HiFi Cas9 Nuclease v3 in 1× CutSmart Buffer for the generation of Ribonucleoprotein (RNP) complexes (all components from Integrated DNA Technologies) according to the ONT protocol (version ENR_9084_v109_revD_04Dec2018).

The dephosphorylation, Cas9-mediated digestion and dA-tailing of 1–10 µg input genomic DNA was carried out according to the ONT protocol. The genomic DNA was incubated with the RNPs for 20 min at 37 °C and then 2 min at 80 °C for enzyme inactivation. ONT sequencing adapters (AMX) were ligated to the cleaved and dA-tailed target ends for 10 min at room temperature before stopping the reaction by adding one volume of 10 mM Tris-EDTA (pH 8.0). Short fragments (< 3 kbp) and residual enzymes were removed by adding 0.3× AMPure XP beads (Beckman-Coulter, Brea, California) and washing twice in long fragment buffer (ONT). The DNA was eluted by incubating for 10 min at room temperature in elution buffer (ONT). Cas9-multiplexing experiments were carried out as described above but EXP-NBD104 native barcodes (ONT) were ligated to the cleaved and dA-tailed target ends using Blunt/TA Ligase Master Mix (New England Biolabs). Samples were quantified and all available nanograms were pooled in a final volume of 65 µL nuclease-free water. ONT sequencing adapters (AMXII from EXP-NBD104) were ligated, followed by purification, washing and elution as described above for the singleplex experiments. The purified library was mixed with 37.5 µL sequencing buffer (ONT) and 25.5 µL of library loading beads (ONT). The library was loaded onto a FLO-MIN106D (R9.4.1) flow cell and sequenced using MinKNOW v20.06.5 (ONT) until a plateau was reached.

### ONT sequence data analysis

Raw fast5 files were base called using Guppy v3.4.5 in high-accuracy mode, with parameters “-r - i $FAST5_DIR -s $BASECALLING_DIR --flowcell FLO-MIN106 --kit SQK-LSK109 -- pt_scaling TRUE” (the last parameter was recommended by ONT technical support to achieve the most appropriate scaling of reads with biased sequence composition, as expected for repeat motifs with low complexity). Reads from multiplexed runs were demultiplexed using Guppy v3.4.5 with parameters “-i $BASECALLING_DIR -s $DEMulTIPLEXING_DIR --trim_barcodes -- barcode_kits EXP-NBD104”. Quality filtering was carried out using NanoFilt v2.7.1(De Coster et al., n.d.), requiring a minimum quality score of 7. Reads spanning the full repeat were identified using a combination of the scripts msa.sh and cutprimers.sh from BBMap suite v38.87 (https://sourceforge.net/projects/bbmap/). In particular, we used *in silico* PCR with 100-bp primers annealing to the flanking regions (minimum alignment identity = 80%). These “complete sequences” were extracted, aligned to either the normal or expanded allele based on length, and subsequent analysis was carried out using a *k*-means clustering method (*k* = 2). Reads from each allele and each sample were then processed separately. Length variability of normal and expanded alleles was determined using the coefficient of variation, measured by the ratio of the standard deviations to the average of the normal/expanded allele lengths.

An accurate consensus sequence was generated by collapsing reads from the normal allele as previously described, using the medaka “r941_min_high_g344” model(Grosso et al. 2021). Finally, the polished consensus sequence was screened for repeats using Tandem Repeat Finder v4.09(Benson 1999). Scripts for the generation of consensus sequences and repeat annotations for the normal allele have been deposited online (https://github.com/MaestSi/CharONT).

Reads from the expanded allele were aligned to sequences flanking the repeat, screened for repeats containing the motifs “TG”, “CCTG” and/or “TCTG”, and visualized in a genome browser(Grosso et al. 2021). The extracted sequences and repeat summary file were then imported into R and used to generate a simplified version of the reads with a custom script. In particular, read coordinates corresponding to annotated repeats were replaced with a single-nucleotide stretch equal in length to the annotated repeat, with each motif corresponding to a different nucleotide stretch. In more detail, the flanking region was reported as unchanged to simplify the alignment of reads, “TG” was replaced with “GG”, “CCTG” with “CCCC”, “TCTG” with “TTTT, and remaining non-annotated portions were replaced with stretches of “N” of the same length. Reverse complements were generated for simplified reads originally in the reverse orientation, and all simplified reads were aligned to the flanking sequence using Minimap2 v2.17-r941 and visualized in Integrative Genomics Viewer (IGV) v2.8.3. Scripts for the annotation of repeats and the generation of simplified reads for the expanded allele have been deposited online (https://github.com/MaestSi/MosaicViewer_CNBP).

## Data Availability

The sequencing data generated in this study have been submitted to the NCBI BioProject database (https://www.ncbi.nlm.nih.gov/bioproject/) under accession number PRJNA818354. For revision purposes, reviewers can download data from the following link: https://dataview.ncbi.nlm.nih.gov/object/PRJNA818354?reviewer=1g4lp9ijkgv5s07is6imnot6ch

## Author Contributions

Conceptualization, M.R., M.D. and A.B.; Methodology, M.A., M.R., S.M. and M.D., Software, S.M.; Validation F.C. and V.V.V.; Formal Analysis, C.D.E. and M.R.D.; Investigation, M.A. and L.D.A.; Resources, R.M.; Writing – original draft presentation, M.A., M.R. and S.M.; Writing – review and editing, A.B. and M.R.; Supervision, G.N., A.B. and M.D.; Funding Acquisition, M.R., M.D. and A.B.

## Acknowledgements

This research was funded by the Muscular Dystrophy Association (www.mda.org). We also thank the Italian DiMio onlus association of DM patients (www.dimio.it) for funding to support research projects.

