## Supplementary for "Characterization of full-length *CNBP* expanded alleles in myotonic dystrophy type 2 patients by Cas9-mediated enrichment and nanopore sequencing"

| Run ID | Exp1_Singleplex | Exp2_Singleplex | Exp3_Singleplex | Exp4_Singleplex | Exp1_Multiplex | Exp2_Multiplex | Exp3_Multiplex | Exp4_Multiplex |
|---|---|---|---|---|---|---|---|---|
| **Source** | Whole blood | Whole blood | Whole blood | Whole blood | Whole blood | Whole blood | Whole blood | Whole blood |
| **# samples** | 1 | 1 | 1 | 1 | 5 | 4 | 4 | 3 |
| **Sample ID** | A2 | E | F | B | A1, A3, A4, C | A1, A3, A4, C | A4, B, D | B, D, F |
| **Input** | 5 ug | 2 ug | 6,8 | 4,6 | 2 ug / sample | 1 - 4 ug / sample | 3 - 5 ug / sample | 4.5 - 10 ug / sample |
| **Total reads** | 186.622 | 40.122 | 27.693 | 92.124 | 1.504.794 | 917.260 | 114.967 | 653.967 |
| **Total aligned PASS reads** | 154.046 | 35.025 | 21.550 | 60.315 | 1.284.125 | 819.341 | 98.083 | 552.544 |
| **On-target PASS reads On CNBP** | 624 | 283 | 330 | 142 | 417 | 397 | 75 | 501 |
| **On-target reads %** | 0,41% | 0,81% | 1,53% | 0,24% | 0,03% | 0,05% | 0,08% | 0,09% |
| **On-target avg. cov. (X)** | 584,2 | 279,8 | 311,9 | 127,6 | 507,81 | 357,54 | 78,35 | 494,44 |
| **Whole genome avg. cov. (X)** | 0,14 | 0,077 | 0,07 | 0,07 | 0,94 | 0,6 | 0,118 | 0,66 |
| **Fold enrichment** | 4.173 | 3.634 | 4.456 | 1.823 | 540 | 596 | 664 | 749 |

| | **Average values** | |
|---|---|---|
| | **Singleplex (N=4)** | **Multiplex (N=4)** |
| **Total aligned reads** | 67734 | 688523 |
| **On-target reads** | 345 | 348 |
| **On-target %** | 0,01 | 0,001 |
| **On-target avg. cov. (X)** | 325,9 | 359,5 |
| **Whole genome avg. cov. (X)** | 0,1 | 0,6 |
| **Fold enrichment** | 3521,3 | 637,3 |

**Table S1. Sequencing statistics of singleplex and multiplex experiments**

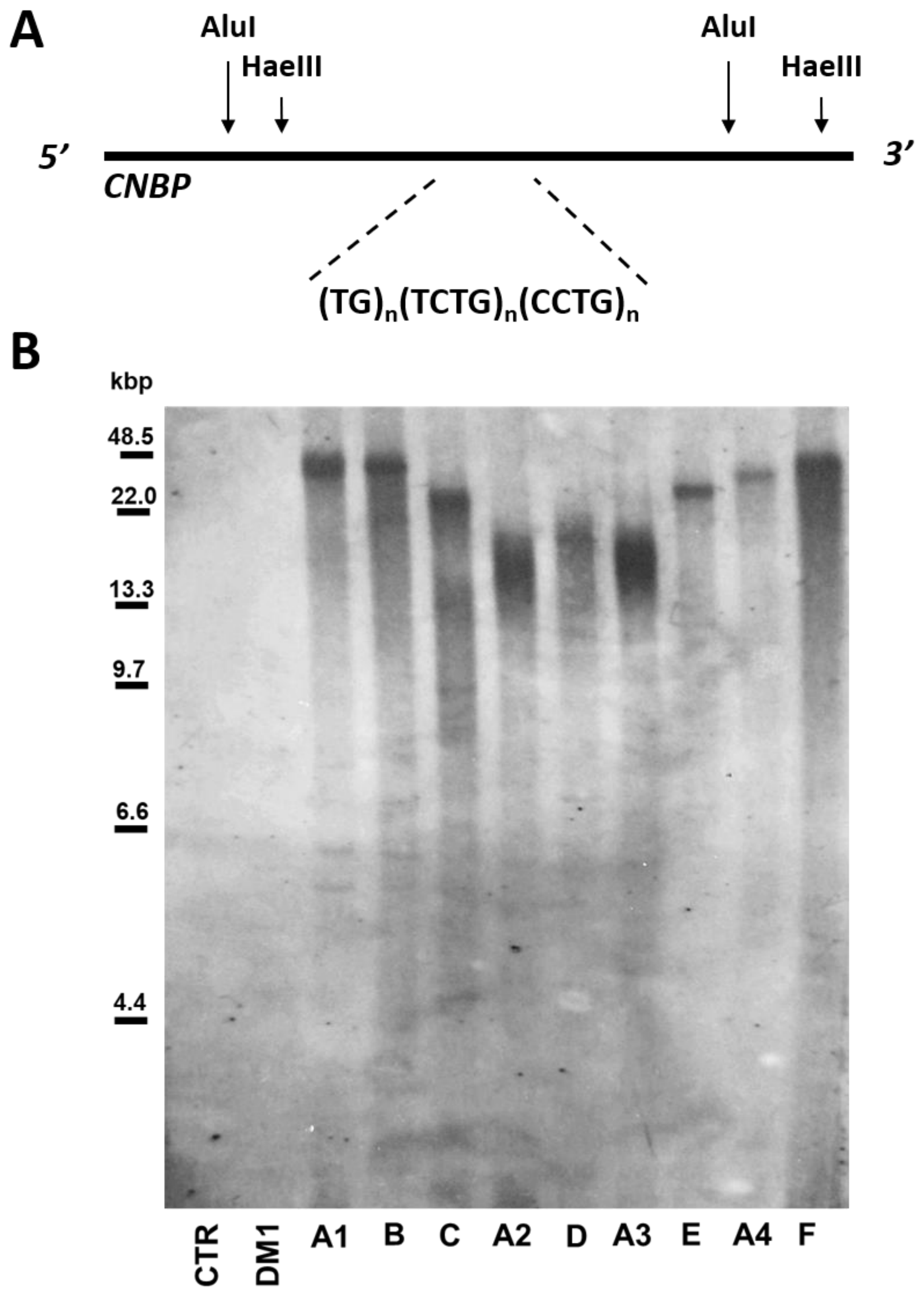


**Figure S1. Southern blot analysis of expanded alleles in DM2 patients. (A)** Restriction map of the *DM2* locus indicating the AluI and HaeIII restriction sites used for the digestion of genomic DNA. **(B)** Southern blot analysis of genomic DNA double digested with AluI and HaeIII and probed with a digoxigenin (DIG)-labeled $(CCTG)_5$ locked nucleic acid (LNA) probe. Lane 1, CTR, healthy control sample; lane 2, DM1 sample; lanes 3–11, DM2 samples. Molecular markers are indicated on the left.

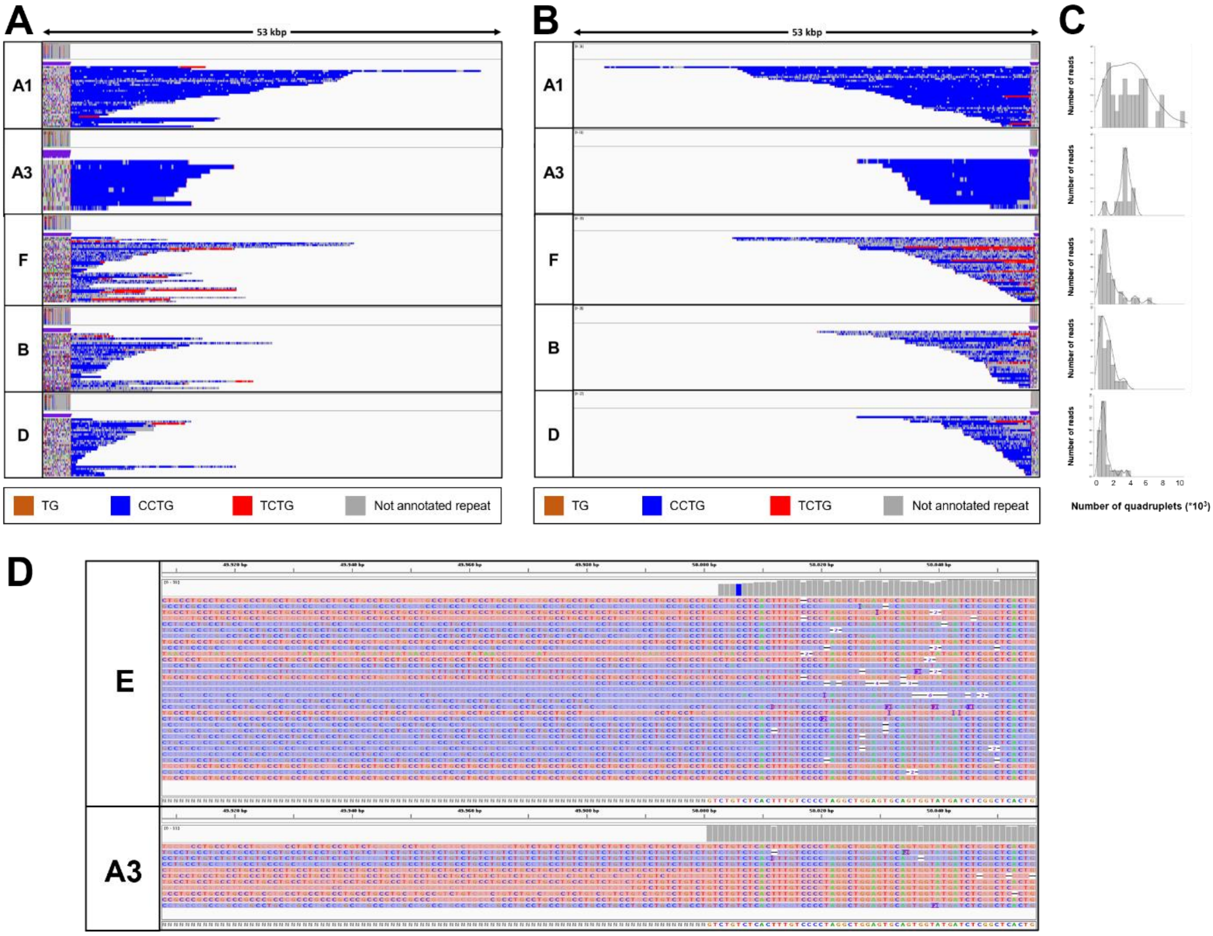


**Figure S2. Analysis of the *CNBP* repeat motif for the expanded alleles in DM2 patients.** Integrative Genomics viewer (IGV) visualization (53-kbp windows) of ONT targeted sequencing data from the expanded alleles of the four remaining DM2 samples. Complete reads were aligned at the 3′ end **(A)** and subsequently at the 5′ end **(B)** in order to identify the repeat pattern characterizing the expanded microsatellite locus. Each motif in the expanded alleles was visualized using a different color, as indicated in the key. All samples feature the unexpected TCTG motif of variable length downstream of the CCTG motif. **(C)** Abundance of quadruplets identified in each patient. The *y*-axis shows the number of ONT reads with a certain number of repeats, whereas the *x*-axis shows the number of quadruplet repeats identified. ONT reads were grouped into 500-bp bins. The gray line represents the estimated kernel density of the underlying solid gray distribution of ONT reads. **(D)** IGV visualization of ONT targeted sequencing data from the expanded alleles of two representative patients (E and A3) carrying a pure CCTG expansion and a repeat with the TCTG motif, respectively.

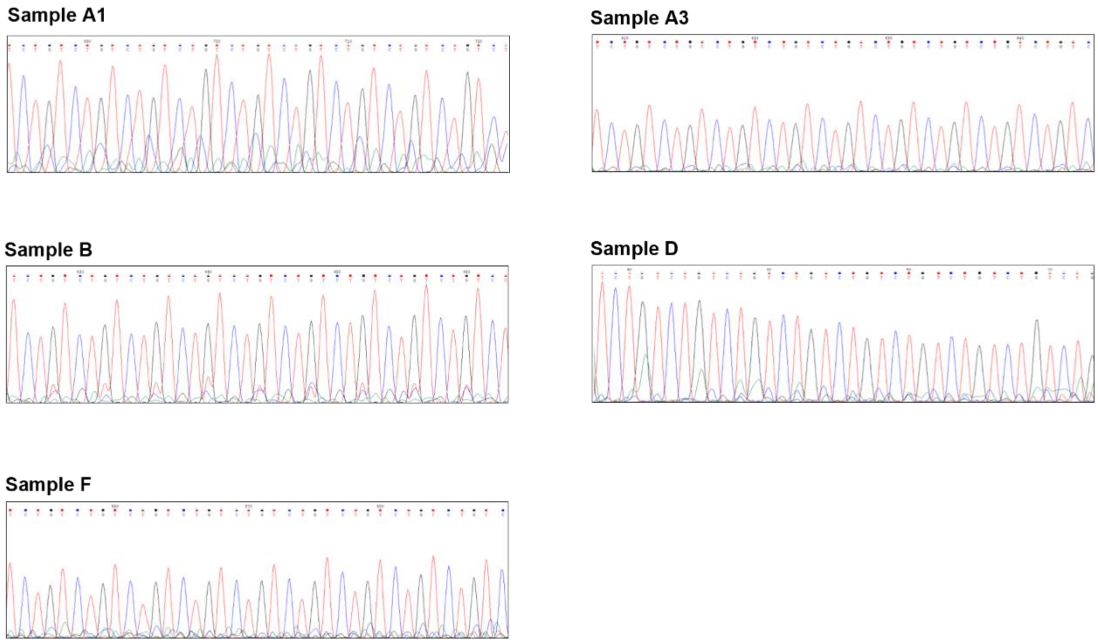


**Figure S3 Sanger sequencing of QP-PCR products showing the presence of the (TCTG)n motif.** QP-PCR was carried out using primer P4TCTG and genomic DNA from DM2 patients A1, A3, B, D and F. Sanger sequencing of the QP-PCR products confirmed the presence of the (TCTG)n array at the 3’ end of the (CCTG) expansions.
